# Lymphocytes contribute to DUX4 target genes in FSHD muscle biopsies

**DOI:** 10.1101/717652

**Authors:** Christopher R. S. Banerji, Maryna Panamarova, Peter S. Zammit

**Affiliations:** King’s College London, Randall Centre for Cell and Molecular Biophysics, New Hunt’s House, Guy’s Campus, London SE1 1UL, UK; Faculty of Medicine, Imperial College London, Level 2, Faculty Building, South Kensington Campus, London SW7 2AZ, UK

**Author notes:** Corresponding Author: Christopher R.S. Banerji, and.

**Keywords:** DUX4, FSHD, facioscapulohumeral muscular dystrophy, lymphocyte, skeletal muscle

## Abstract

Facioscapulohumeral muscular dystrophy (FSHD) is an incurable myopathy linked to overexpression of DUX4. However, DUX4 is difficult to detect in FSHD myoblasts and target gene expression is not a consistent FSHD muscle biopsy biomarker, displaying efficacy only on pathologically inflamed samples. Immune gene misregulation occurs in FSHD muscle biopsies with DUX4 targets enriched for inflammatory processes. However, assessment of the FSHD immune cell transcriptome, and the evaluation of DUX4 and target gene expression has not yet been performed. We show that FSHD lymphoblastoid cell lines (LCLs) display robust DUX4 expression, and express early and late DUX4 targets. Moreover, genes elevated on FSHD LCLs are elevated in FSHD muscle biopsies, correlating with DUX4 target activation and histological inflammation. These genes are importantly unaltered in FSHD myoblasts/myotubes, implying a non-muscle source in biopsies. Our results indicate an immune cell source of DUX4 and target gene expression in FSHD muscle biopsies.

## Introduction

Facioscapulohumeral muscular dystrophy (FSHD) is a prevalent (12/100,000^1^) inherited skeletal myopathy. Clinically, FSHD manifests as a descending skeletal muscle atrophy, commencing in the facial muscles before progressing to the shoulder girdle, associating with scapular winging, and latterly the muscles of the lower limb, resulting in foot drop^2,3^. The pattern of muscle involvement in FSHD is characteristically left/right asymmetric^4^. Heterogeneity in pathology progression among first degree relatives, including monozygotic twins, is also well described^5–7^. Moreover, systemic disruption in FSHD pathology is suggested by several extra-muscular features including retinal telangiectasia similar to Coat’s disease^8–10^ and sensorineural hearing loss^11,12^.

FSHD shows an autosomal dominant pattern of inheritance linked to hypomethylation of the D4Z4 macro-satellite at chromosome 4q35^2^. This epigenetic modification can be achieved in two ways: ∼95% of cases (FSHD1 - MIM 158900) have truncation of the D4Z4 region to 1-10 repeats^13^, while the remaining ∼5% (FSHD2 - MIM 158901) carry mutations in chromatin modifying genes such as *SMCHD1*^14^, or more rarely, *DNMT3B*^15^. In addition to D4Z4 hypomethylation, FSHD patients also carry a permissive 4qA haplotype, encoding a poly(A) signal^13^. Each 3.3kb D4Z4 repeat unit encodes an open reading frame for the transcription factor double homeobox 4 (DUX4). Epigenetic derepression of D4Z4 via hypomethylation permits expression of DUX4 transcripts from the most distal repeat, which are then stabilised by the poly(A) signal for translation. Mis-expression of DUX4 protein is thus proposed to underlie FSHD pathology.

How DUX4 drives pathology in FSHD is poorly understood. DUX4 expression in myoblasts has been shown to induce a set of genes which are pro-apoptotic^16–18^, anti-myogenic^18,19^ and curiously, immune system related^20,21^. However, DUX4 detection in FSHD patient derived myogenic cultures is notoriously difficult, with expression reported to be as low as 1/1000-1/5000 cells in myoblasts and 1/200 myonuclei in differentiated myotubes^20,22^. DUX4 target gene expression has been proposed as a biomarker of FSHD muscle biopsies^21^, but we have demonstrated via meta-analysis that its discriminatory power is generally underwhelming: appreciable levels of DUX4 target genes are detectable only in muscle biopsies which have been preselected for active disease/inflammation, via MRI metrics of T1 and STIR positivity^23,24^. Given this disappointing result, we investigated other biomarkers for FSHD muscle biopsies. DUX4 shows homology with the homeodomains of the myogenic master regulator PAX7 and a competitive interaction has been proposed between the two proteins^19,25^, with PAX7 homeodomains able to substitute those of DUX4 without affecting certain DUX4 functions^25^. We demonstrated that a biomarker based on suppression of PAX7 target genes hallmarks FSHD muscle biopsies, as well as isolated myoblasts, significantly outperforming DUX4 target gene expression^23,24^.

PAX7 target gene repression and DUX4 target gene activation are however, independently associated with the degree of histological inflammation and active disease in a recent MRI guided FSHD muscle biopsy study^26^, implying that both mechanisms contribute to pathology^24^. Given that DUX4 is expressed at such low levels in patient muscle cells, the question remains as to which cells are expressing DUX4 and its target genes in these highly inflamed biopsies? Histological evidence of muscle inflammation in FSHD is well documented^27–30^, with perivascular and endomysial lymphocytic (predominantly CD8^+^) infiltrates a consistent finding, which is most robust on STIR positive muscle biopsies. Elevated circulating levels of pro-inflammatory cytokines such as TNFα have also been reported as inversely associated with maximal voluntary contraction of quadriceps in FSHD^31^.

DUX4 can induce expression of immune system-related genes in myoblasts^20^, and inflammatory genes are dysregulated in FSHD muscle biopsies^21,32^. Recently a large library of 114 control and FSHD lymphoblastoid cell lines (LCLs) (EBV immortalised B-lymphocytes) from 12 FSHD1 affected families was published^33^. In contrast to studies of FSHD myoblasts all 61 FSHD LCLs displayed robust DUX4 expression, as well as DUX4 target genes (although only a few were evaluated). Curiously, a small number of control LCLs also express DUX4, albeit at significantly lower levels to FSHD LCLs^33^. Of further relevance, an in frame DUX4-IGH fusion gene in which the homeodomains of DUX4 are fused to a clamp-like transactivation domain of IGH, presents in a significant subset of B-cell Acute Lymphoblastic Leukaemia (B-ALL). DUX4-IGH can inhibit terminal differentiation and induce transformation^34,35^.

DUX4 expression in FSHD patient derived immune cells may thus represent a non-muscle contributor to pathology, and associate with the elevated levels of DUX4 target genes in inflamed FSHD muscle biopsies. To our knowledge, this has not been directly assessed.

Here we performed RNA-seq of FSHD and control LCLs and primary myoblasts and myotubes to analyse DUX4 and its target gene expression, and to generate an FSHD lymphoblast biomarker. All 3 FSHD LCL lines expressed DUX4 on RNA-seq, compared with no detectible DUX4 in 18 FSHD myoblast samples and DUX4 detected in only 3/18 FSHD myotube samples. We next evaluated expression of both early (8 hours) and late (24-28 hours) DUX4 target genes by assessment of 3 independent DUX4 target gene sets^23^. FSHD LCLs had high expression of both early and late DUX4 target genes in a manner that correlates with DUX4 expression. However, FSHD myoblasts only expressed late DUX4 target genes (implying historic expression of DUX4), while FSHD myotubes had expression of both early and late DUX4 target genes, but in a manner uncorrelated with DUX4 expression (implying a transient DUX4 pulse during differentiation). Thus FSHD LCLs display a more robust expression of DUX4 and DUX4 target genes than FSHD myoblasts or myotubes.

We next performed differential expression analysis of FSHD LCLs compared to controls, deriving an FSHD lymphoblast biomarker of 237 upregulated genes in FSHD LCLs. Crucially, this FSHD lymphoblast biomarker was unaltered between FSHD and control myoblasts or myotube samples, implying that it is not associated with FSHD muscle cells. Performing meta-analysis over transcriptome studies of 7 independent FSHD muscle biopsy datasets describing 130 FSHD samples and 98 controls, we found significant up-regulation of our FSHD lymphoblast biomarker: which was significantly correlated with expression of DUX4 target genes.

Finally, we evaluated the correlation between our FSHD lymphoblast biomarker and histological and MRI assessments of FSHD disease activity and inflammation in muscle biopsies. Using a multivariate model, the FSHD lymphoblast biomarker associated specifically with microscopic histological inflammation, while late DUX4 target gene expression associated with macroscopic MRI based STIR positive inflammation. This implies that our FSHD lymphoblast biomarker is detecting early microscopic inflammatory infiltrates in FSHD muscle before widespread pathological damage is clear on MRI.

In summary, we postulate a role for DUX4 in immune cells in FSHD, and propose that DUX4 target gene expression in FSHD muscle biopsies may not solely be driven by muscle cells. DUX4 target gene expression in immune cells is likely to contribute to FSHD pathology, being associated with early inflammatory changes, at a time when therapeutic intervention may prevent irreversible change.

## Results

### FSHD LCLs display robust DUX4 expression

We identified 3 FSHD1 patients with gender and first degree relative matched controls, which displayed large reported differences in DUX4 expression from the LCL cohort described by Jones et al., 2017^33^. Namely: GSM16283 (FSHD1, 6RU, female, family 2) and matched control GSM16281; GSM16414 (FSHD1, 6RU, female, family 11) and matched control GSM16320; GSM16278 (FSHD1, 6RU, male, family 2) and matched control GSM16412. RNA-seq was performed on each cell line in triplicate.

We also performed RNA-seq on 3 primary FSHD myoblast cell lines described previously^16^, namely FSHD3 (FSHD1, 7RU, female), FSHD6 (FSHD1, 8RU, female) and FSHD9 (FSHD1, 7RU, male) alongside age and gender matched controls, in proliferation and after 3 days of differentiation into multinucleated myotubes, in singlet. This new RNA-Seq data was considered with our previously published datasets of immortalised FSHD myoblasts and myotubes in triplicate^23,36^. This data describes 3 pathological FSHD cell lines (54-12, 54-A5 and 54-2, all FSHD1, 3RU, male) alongside 2 control lines (54-A10, 54-6, 11RU) from a mosaic patient^37^ as well as 2 further FSHD cell lines (16Abic, FSHD1, 7RU, female and 12Abic, FSHD1, 6RU, female) alongside sibling and gender matched controls (16Ubic and 12Ubic respectively). Thus, a total of 27 immortalised myoblasts and 27 immortalised myotube RNA-seq samples.

DUX4 transcripts were detected by RNA-seq in all FSHD LCL samples (9/9, 100%), and in 2/3 replicates of control LCL GSM16320 (2/9, 22%), although at significantly lower levels than its matched FSHD LCL GSM16414 (Fig. 1A). After adjusting for gender and patient-control pair we found that DUX4 expression was significantly higher in FSHD LCLs compared to controls (*p=*0.0099).

**Figure 1:**
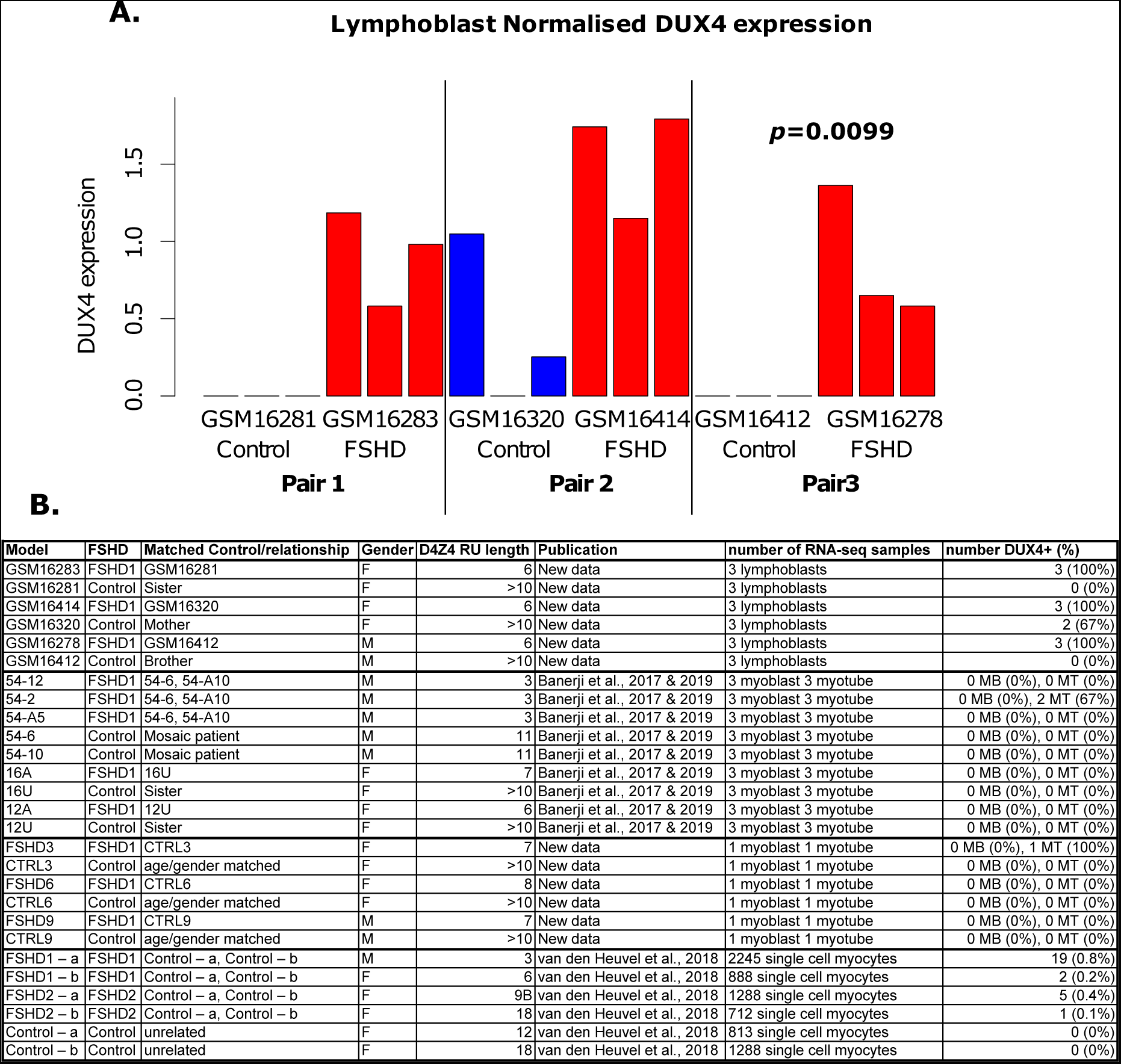
DUX4 expression is robustly detected in RNA-seq of FSHD LCLs. A bar plot displays normalised DUX4 expression for each sample of our RNA-seq of 3 FSHD LCLs and first degree relative matched controls profiled in triplicate (A). The *p*-value denotes the significance of differential expression analysis performed using the DESeq2 package in R, after adjustment for gender and matched pair. A table summarises DUX4 expression in RNA-seq data corresponding to FSHD cellular models (B). Myoblasts and in vitro differentiated myotube data are either new data (primary cell lines) or data previously published by ourselves^23,36^. Single cell RNA-seq of FSHD and control myocytes was previously published by van den Heuvel et al., 2019^32^. A sample was assessed as DUX4 positive if a single detectable DUX4 read was found in normalised RNA-seq data.

In contrast, none of the FSHD or control, primary or immortalised myoblast samples expressed a single DUX4 transcript detectable by RNA-seq (0/18, 0% FSHD; 0/15, 0% controls, Fig. 1B). Considering myotube transcriptomes, 3 FSHD samples expressed detectible DUX4 (3/18, 17%, Fig. 1B), namely primary line FSHD3 and 2/3 replicates of the immortalised 54-2 FSHD cell line. No control myotube samples expressed DUX4 (0/15, 0%, Fig. 1B). In a recent single cell RNA-seq of combined FSHD1 and FSHD2 unfused myocytes, we found DUX4 expression in 27/5133 (0.5%) FSHD cells^32^.

### FSHD LCLs and FSHD myotubes express high levels of early and late DUX4 target genes while FSHD myoblasts express only late DUX4 target genes

We next considered expression of DUX4 target genes in our LCL, myoblast and myotube transcriptomic analysis. We previously described 2 DUX4 target gene expression signatures derived from transcriptomic assessments of human myoblasts over-expressing DUX4 for different lengths of time^23^. A set of 212 DUX4 target genes were derived from data described by Choi et al., 2016^38^ in which DUX4 expression was induced in a genetically modified control myoblast line for 8 hours before samples were collected in triplicate for RNA-seq alongside uninduced controls. Thus, the Choi et al., 2016 DUX4 target gene expression signature represents early DUX4 target genes. A set of 165 DUX4 target genes were derived from data described by Geng et al., 2012^20^, in which control myoblasts were transduced by either a DUX4-encoding, or control, lentiviral vector and samples collected in quadruplicate 24 hours later for microarray analysis. Thus, the Geng et al., 2012 DUX4 target gene expression signature represents later DUX4 target genes.

A further 114 DUX4 target gene signature was described by Yao et al., 2014^21^. RNA-seq data used to derive this signature corresponds to two different control myoblasts: 54-1 transfected with DUX4 lentivirus for 48 hours, and MB135 transfected with DUX4 lentivirus for 24 hours, alongside 54-1 un-transfected control (with contaminating reads from a DUX4 expressing sample) and MB135 transfected with GFP lentivirus for 24 hours^39^. To provide a point of comparison, given its prolific use in recent publications on FSHD, we also consider the Yao et al., 2014 signature as later DUX4 target genes, alongside the Geng et al., 2012 signature.

For DUX4, and each of the 3 DUX4 target gene expression signatures, we computed the mean expression of the genes in each of the LCL, myoblast and myotube samples, to generate a single sample score, as previously described^23,24^. Scores were then *z*-normalised within patient-matched control groups and their performances as biomarkers of FSHD status evaluated using ROC curves, separately for LCLs, myoblasts and myotubes (Fig. 2).

**Figure 2:**
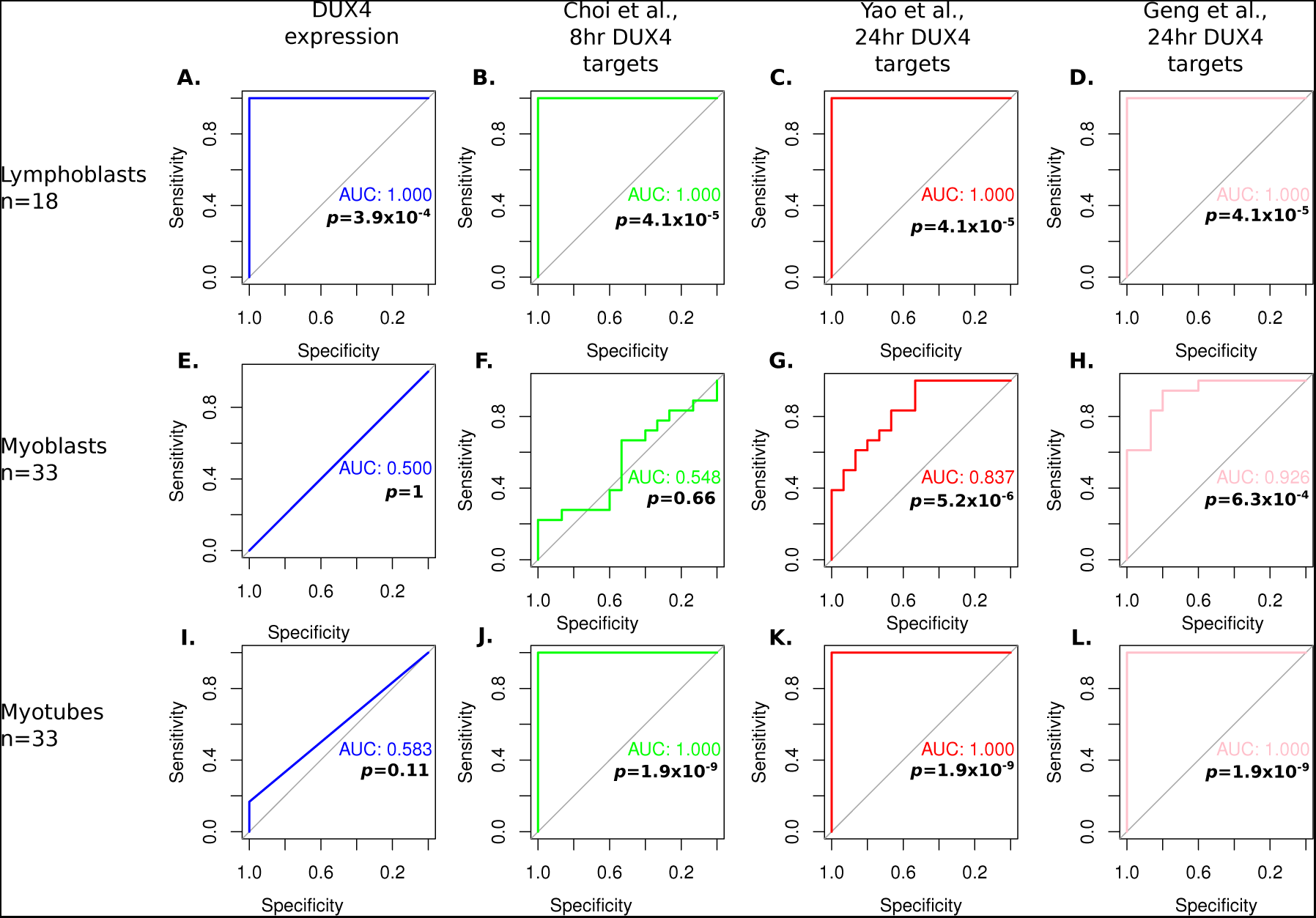
DUX4 and early and late DUX4 target gene expression identifies FSHD LCLs more robustly than FSHD myoblasts or myotubes. ROC curves display the discriminatory power of DUX4 expression or expression of DUX4 target genes in patient derived LCLs (A-D), myoblasts (E-H) and in vitro differentiated myotubes (I-L), using the early Choi et al., 2016^38^ (8 hour) DUX4 target gene signature, and the later Yao et al., 2014^21^ (24-48 hour) and Geng et al., 2012^20^ (24 hour) DUX4 target gene signatures (all *z-*normalised within FSHD patient matched control group within cell type). AUC for each discriminator in each cell line is displayed alongside Wilcoxon *p*-values comparing the normalised biomarker value in FSHD samples vs controls. Only on LCLs are all 4 biomarkers perfect discriminators of FSHD status.

For LCLs, DUX4 expression, and each of the 3 DUX4 target gene expression signatures were perfect classifiers of FSHD status (Wilcoxon *p*<3.9×10^−4^, AUC=1, n=18 (9 FSHD, 9 control), Fig. 2A-D). For myoblasts, no sample expressed DUX4, and the Choi et al., 2016 early DUX4 target gene expression signature did not represent a significant classifier of FSHD status (FSHD vs control: Wilcoxon *p*=0.66, AUC=0.548, n=33 (18 FSHD, 15 control), Fig. 2E-F). However, both the late DUX4 target gene signatures were significant classifiers of FSHD myoblasts (Geng et al., 2012, Wilcoxon *p*=5.2×10^−6^, AUC=0.926, Yao et al., 2014 Wilcoxon *p*=6.3×10^−4^ AUC=0.837, Fig. 2G-H). Therefore, although FSHD myoblasts do not express DUX4, nor hold the hallmarks of recent DUX4 target gene expression, they do express late DUX4 target genes, implying historic DUX4 expression. For myotubes, DUX4 expression did not represent a significant classifier of FSHD status (FSHD vs control: Wilcoxon *p*=0.11, AUC=0.583, n=33 (18 FSHD, 15 control), Fig. 2I). However, both the early and the 2 late DUX4 target gene expression signatures were perfect classifiers of FSHD myotubes (Wilcoxon *p*=1.9×10^−9^, AUC=1, Fig 2J-L). This suggests that during myogenic differentiation, FSHD myoblasts express a transient pulse of DUX4 expression, leading to activation of both early and late DUX4 targets by the end of differentiation, although DUX4 itself has been suppressed by this point.

### DUX4 expression is correlated with early and late DUX4 target gene expression in FSHD LCLs but not in FSHD myotubes

We next investigated how DUX4 and DUX4 target genes correlated with one another within the different cell types. For LCLs, DUX4 expression correlated strongly with both early and late DUX4 target gene expression (DUX4 expression vs Choi et al., *p*=5.3×10^−5^, Pearson’s *r=*0.81, DUX4 expression vs Geng et al., *p*=5.3×10^−4^, Pearson’s *r=*0.78, DUX4 expression vs Yao et al., *p*=1.5×10^−5^, Pearson’s *r=*0.78, Fig. 3A). The early and late DUX4 target gene expression scores also correlated strongly in LCLs (Choi et al., vs Geng et al., *p*=1.2×10^−10^, Pearson’s *r=*0.96, Choi et al., vs Yao et al., *p*=8.7×10^−10^, Pearson’s *r=*0.95, Fig. 3A). This confirms that the DUX4 target genes identified via exogenous DUX4 expression in myoblasts, associates with endogenous DUX4 expression in FSHD LCLs, implying a cell type ubiquity of DUX4 target gene activation. This suggests that DUX4 target genes detected in FSHD muscle biopsies may be derived from immune cells, as well as muscle cells.

**Figure 3:**
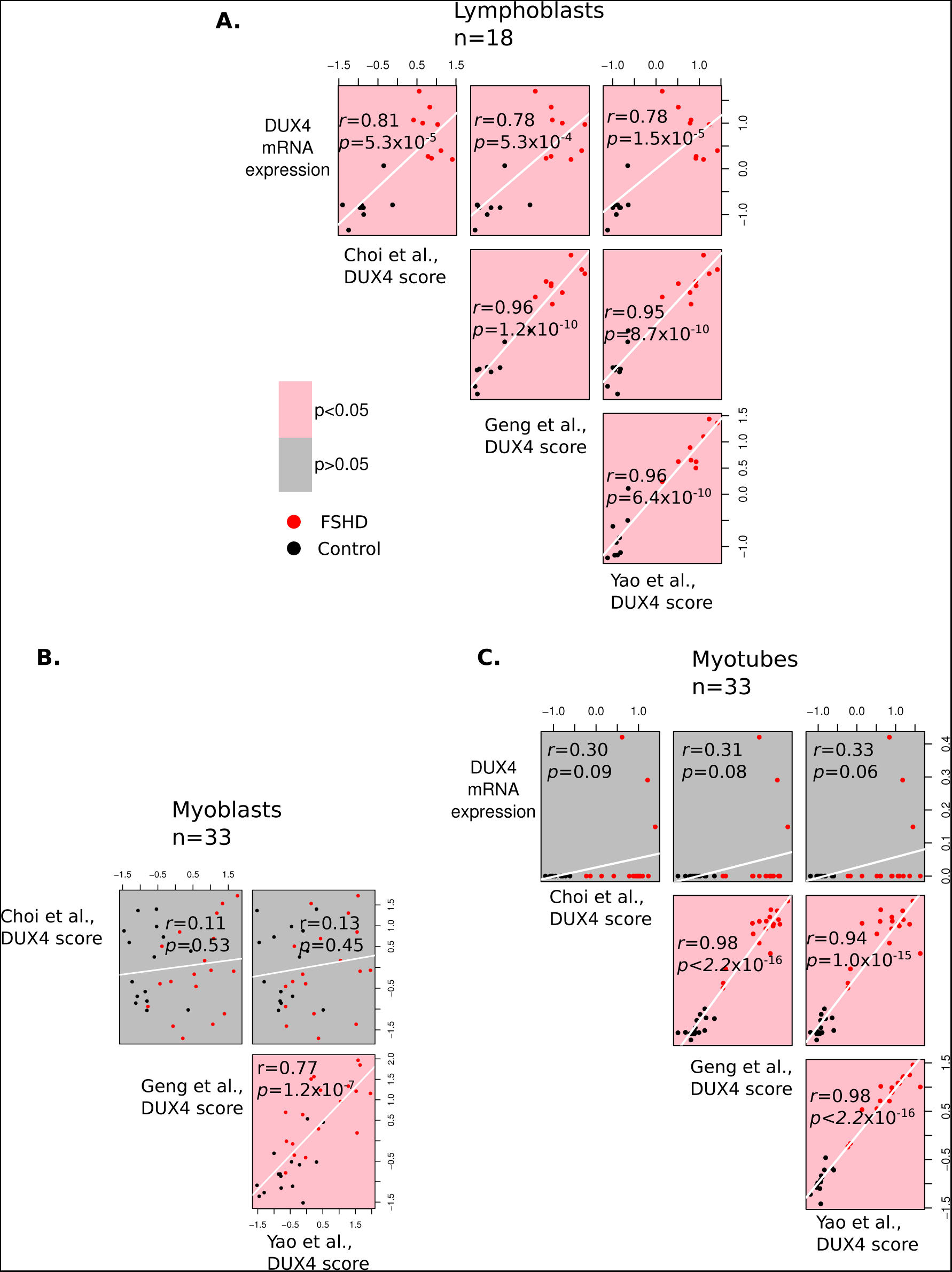
DUX4 expression correlates with expression of early and late DUX4 target genes in LCLs but not in myoblasts or myotubes. Scatter plots display DUX4 expression, and DUX4 target genes using the early Choi et al., 2016^38^ (8 hour), and the later Yao et al., 2014^21^ (24-48 hour) and Geng et al., 2012^20^ (24 hour) DUX4 target gene signatures (all *z-*normalised within FSHD patient matched control group within cell type) plotted against one another across the 18 LCL samples (A), the 33 myoblast samples (B) and the 33 myotube samples (C). Pearson’s *r* and associated *p*-value are provided for each pairwise comparison. Red points correspond to FSHD samples, while black points correspond to controls. Plots denoting correlations reaching significance are pink, whilst those not attaining significance are grey. Since no myoblast samples expressed DUX4, the DUX4 mRNA expression comparison row is not displayed (B). Only on LCLs are all 4 DUX4 biomarkers significantly correlate

In myoblasts, which all lacked DUX4 expression, the two late DUX4 target gene signatures of Geng et al., 2012 and Yao et al., 2014 correlated (Geng et al., vs Yao et al., *p*=1.2×10^−7^, Pearson’s *r=*0.77, Fig. 3B), confirming their reproducibility, while the Choi et al., 2016 early DUX4 target gene expression signature was unrelated to these later DUX4 target gene sets (Choi et al., vs Geng et al., *p*=0.45, Pearson’s *r=*0.11, Choi et al., vs Yao et al., *p*=0.45, Pearson’s *r=*0.13, Fig. 3B). This implies that DUX4 expression in FSHD myoblasts is sufficiently historic to ensure that early DUX4 target gene expression cannot be related to persistent late target gene activation.

In the myotubes we found no association between DUX4 expression and any of the DUX4 target gene scores (DUX4 expression vs Choi et al., *p*=0.09, Pearson’s *r=*0.30, DUX4 expression vs Geng et al., *p*=0.08, Pearson’s *r=*0.31, DUX4 expression vs Yao et al., *p*=0.06, Pearson’s *r=*0.33 Fig. 3C), but this analysis is underpowered as only 3 samples expressed DUX4. In contrast to myoblasts however, there was a strong correlation between the early and late DUX4 target gene expression scores (Choi et al., vs Geng et al., *p<*2.2×10^−16^, Pearson’s *r*=0.98, Choi et al., vs Yao et al., *p*=1.0×10^−15^, Pearson’s *r=*0.94, Fig. 3C). This implies a transient burst of DUX4 expression during differentiation. In line with this, FSHD myotube samples express significantly higher levels of both early and late DUX4 target genes than their corresponding FSHD myoblast samples (Wilcoxon *p<*1.4×10^−4^ Fig S1A-C). Conversely, control myotubes display significantly lower levels of the early DUX4 target genes (Wilcoxon *p=*0.002, FigS1D), and similar levels of late DUX4 targets (Wilcoxon *p*>0.3, FigS1E-F) to their corresponding myoblast samples.

We previously evaluated the discriminatory power of the 3 DUX4 target gene scores on unfused FSHD myocytes profiled by single cell RNA-seq, and although significant discriminators, no score achieved an AUC>0.56^24^. However, 27/5133 myocytes expressed DUX4^32^, which offers greater power for assessment of DUX4 association with the target gene scores in differentiated muscle cells, compared to the 3 DUX4 positive myotube samples. Surprisingly, while the early and late DUX4 target gene expression scores correlated in this single cell data set (Choi et al., vs Geng et al., *p<*2.2×10^−16^, Pearson’s *r*=0.54, Choi et al., vs Yao et al., *p*<2.2×10^−16^, Pearson’s *r=*0.38, Geng et al., vs Yao et al., *p*<2.2×10^−16^, Pearson’s *r=*0.86 Fig. S2), DUX4 expression across DUX4 positive cells was again not associated with either early nor late DUX4 targets in single FSHD myocytes (DUX4 expression vs Choi et al., *p*=0.8, Pearson’s *r=*0.16, DUX4 expression vs Geng et al., *p*=0.6, Pearson’s *r=*0.22, DUX4 expression vs Yao et al., *p*=0.5, Pearson’s *r=*0.23, Fig. S2). Plotting DUX4 expression against early and late DUX4 target gene expression scores in the single cell data, reveals a peak of DUX4 expression in cells with low levels of DUX4 target genes, which decays as DUX4 target genes increase (Fig S2). This is consistent with a transient pulse of DUX4 expression occurring in differentiating FSHD myoblasts, which shuts down as DUX4 targets are activated.

### An FSHD lymphoblast signature is up-regulated in FSHD muscle biopsies and correlates with DUX4 target gene expression

Given that FSHD LCLs express high levels of DUX4 and both early and late DUX4 target genes, and that FSHD muscle biopsies are often characterised by inflammation in a manner correlating with DUX4 target gene expression^24,26^, we next investigated whether an FSHD LCL-derived gene expression signature can discriminate FSHD muscle biopsies from controls.

We performed a differential expression analysis comparing FSHD LCLs to controls, adjusting for gender and sibling matched pairs. We identified a large number of differentially expressed genes and considered the 500 most significantly altered for further analysis. Of these 237 were up-regulated in FSHD lymphocytes. As DUX4 is a known transcriptional activator and genes suppressed under DUX4 expression do not add power to DUX4 target based FSHD biomarkers^23^, we considered the mean expression of these 237 FSHD LCL up-regulated genes in a given sample as a putative FSHD biomarker, hereafter called the FSHD Lymphoblast score (**Table S1**). Of these 237 genes, 9 were also present in the Choi et al. and 1 in the Geng et al. DUX4 target gene signatures, but none in that of Yao et al. The full FSHD Lymphoblast score is used here, however the significance of results are unchanged when these overlapping genes are removed.

The FSHD Lymphoblast score was evaluated on each sample of 7 independent FSHD muscle biopsy transcriptomic studies^21,26,42–46^, totalling 130 FSHD samples alongside 98 matched controls. The FSHD Lymphoblast score was significantly upregulated in FSHD muscle biopsies on meta-analysis (Fisher’s combined *p*=0.0007, Fig. 4A), achieving outright significance on 2/7 data sets, and representing a moderately powered biomarker of FSHD status under ROC curve analysis (Wilcoxon *p*=0.0018, AUC=0.621, Fig. 4B). Of the FSHD muscle biopsy data sets, the strongest upregulation of the FSHD Lymphoblast score was found in the MRI guided RNA-seq dataset^26^, in which all but 2 FSHD samples displayed STIR positivity, indicative of active inflammation (Wang et al., 2019, Wilcoxon *p*<1.5×10^−5^, Fig. 4A).

**Figure 4:**
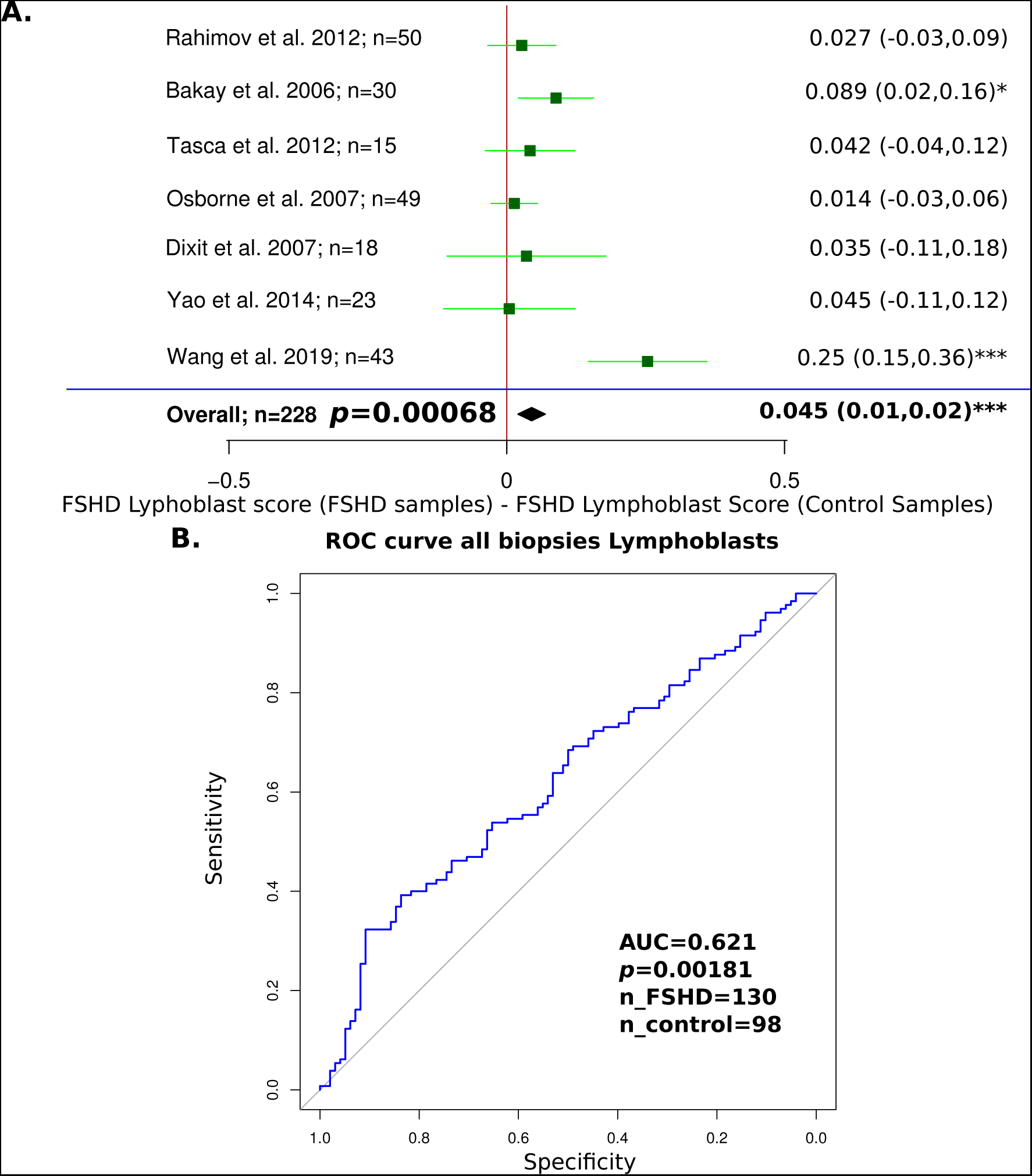
The FSHD Lymphoblast score is elevated on FSHD muscle biopsies compared to controls on meta-analysis of 7 independent data sets. A forest plot displays the significance of the FSHD Lymphoblast score as a discriminator of FSHD muscle biopsies in the 7 independent data sets assessed (A). Green boxes denote the mean difference in FSHD Lymphoblast score between FSHD and control muscle biopsies and whiskers denote the 95% confidence interval. A vertical line denotes a score difference of 0 and datasets where the whiskers cross this line have not attained significance at the 5% level (as assessed by Wilcoxon *U*-test). Numerical values for mean score difference and confidence interval are displayed for each dataset on the left of the plot with significance denoted by asterisks as follows: *: *p*<0.05, **: *p*<0.01, ***: *p*<0.001. The overall estimate is displayed as a diamond and was computed using a random effects model with significance assessed via Fisher’s combined test. On meta-analysis the FSHD Lymphoblast score is elevated on FSHD samples, and achieves strongest significance on the Wang et al., 2019^26^ dataset, where muscles displaying evidence of active disease on MRI were preferentially biopsied. A ROC curve displays the discriminatory capacity of the FSHD Lymphoblast score on all muscle biopsy datasets combined (B). The FSHD Lymphoblast score was computed on each muscle biopsy sample and *z*-normalised within each of the 7 independent studies before being pooled for ROC curve analysis. The AUC of the FSHD Lymphoblast score as a discriminator of FSHD muscle biopsies is displayed alongside the Wilcoxon *p*-value comparing normalised FSHD Lymphoblast score values in FSHD muscle biopsies to controls

Importantly, upregulation of the FSHD Lymphoblast score in FSHD muscle biopsies is unlikely to be driven by muscle gene expression, since there was no significant difference in expression of the FSHD Lymphoblast score on our RNA-seq data of FSHD and control myoblasts (Wilcoxon *p*=0.76, Fig. 5A) and myotubes (Wilcoxon *p*=0.81, Fig. 5B).

**Figure 5:**
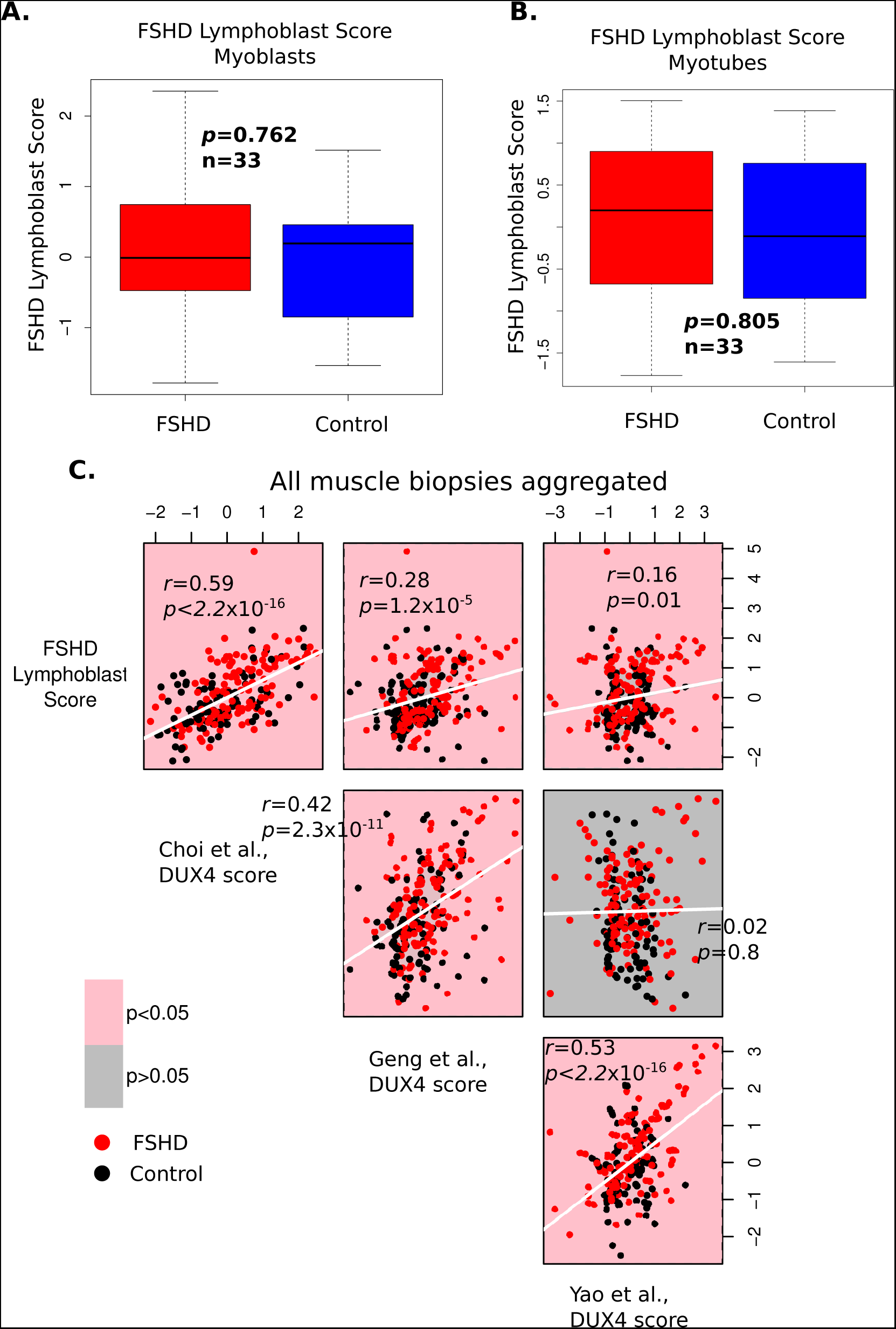
The FSHD Lymphoblast score is unaltered on FSHD myoblast and myotube samples and correlates with the level of DUX4 target gene expression in FSHD muscle biopsies. Box plots display the FSHD Lymphoblast score (*z-*normalised within FSHD patient matched control group within cell type) in FSHD and control myoblast samples (A) and myotube samples (B). The box represents the interquartile range (IQR), with the median indicated by a line. Whiskers denote min (1.5*IQR, max (observed value)). Wilcoxon *U*-test *p*-values comparing FSHD to control samples are presented and demonstrate that the FSHD Lymphoblast score is not significantly altered in FSHD on either myoblasts or myotubes. Scatter plots display the FSHD Lymphoblast score, the early Choi et al., 2016^38^ (8 hour), and later Yao et al., 2014^21^ (24-48 hour) and Geng et al., 2012^20^ (24 hour) DUX4 target gene signatures (all *z-*normalised within each of the 7 muscle biopsy studies) plotted against one another across all 228 muscle biopsies (130 FSHD, 98 control) (C). Pearson’s *r* and associated *p*-value is provided for each pairwise comparison. Red points correspond to FSHD samples, while black points correspond to controls. Plots denoting correlations reaching significance are pink, whilst those not attaining significance are grey. The FSHD Lymphoblast score correlates with all the DUX4 target gene expression scores but most strongly with the early DUX4 target gene signature of Choi et al., 2016^38^.

We next computed expression of the 3 DUX4 target gene scores in the muscle biopsy data sets. In line with our previous results, these formed weak but significant discriminators of FSHD status, statistically equivalent to the FSHD Lymphoblast score, but inferior classifiers of FSHD muscle biopsies to PAX7 target gene repression (Fig. S3). Evaluating associations between the FSHD Lymphoblast score and the 3 DUX4 target gene expression scores across the FSHD muscle biopsies revealed that the FSHD Lymphoblast score strongly associated with the early DUX4 target genes (FSHD Lymphoblast score vs Choi et al., *p*<2.2×10^−16^, Pearson’s *r=*0.59, Fig. 5C). A weaker but significant association was found between the FSHD Lymphoblast score and the two late DUX4 target gene expression signatures (FSHD Lymphoblast score vs Geng et al., *p*=1.5×10^−5^, Pearson’s *r=*0.28, FSHD Lymphoblast score vs Yao et al., *p*=0.01, Pearson’s *r=*0.16 Fig. 5B).

### The FSHD Lymphoblast score is associated with histological inflammation in FSHD muscle biopsies, independently of DUX4 target gene expression

FSHD LCL gene expression is elevated in FSHD muscle biopsies but not FSHD myoblasts or myotubes, suggesting that the FSHD Lymphoblast score may be detecting immune cell infiltrates in FSHD muscle biopsies. To investigate, we considered published RNA-seq data of FSHD muscle biopsies alongside histological assessment of pathology score, inflammation and active disease, together with MRI assessment of STIR and T1 positivity and fat fraction^26^. Histological and MRI assessments are all metrics of active pathology in FSHD and hence cross-correlate. We therefore built multivariate regression models evaluating which of these variables was independently associated with the FSHD Lymphoblast score, or each of the 3 DUX4 target gene expression signatures in turn.

Crucially, the FSHD Lymphoblast score associated only with histological inflammation (*p*=0.016, Fig. 6), indicating that our score does indeed correlate with immune cell infiltration of FSHD muscle biopsies. Early DUX4 target genes (Choi et al.) did not independently associate with any of the measures of active pathology in FSHD, but the two late DUX4 target gene expression signatures both significantly associated with STIR positivity (Geng et al., *p*=0.030, Yao et al., *p*=0.020, Fig. 6).

**Figure 6:**
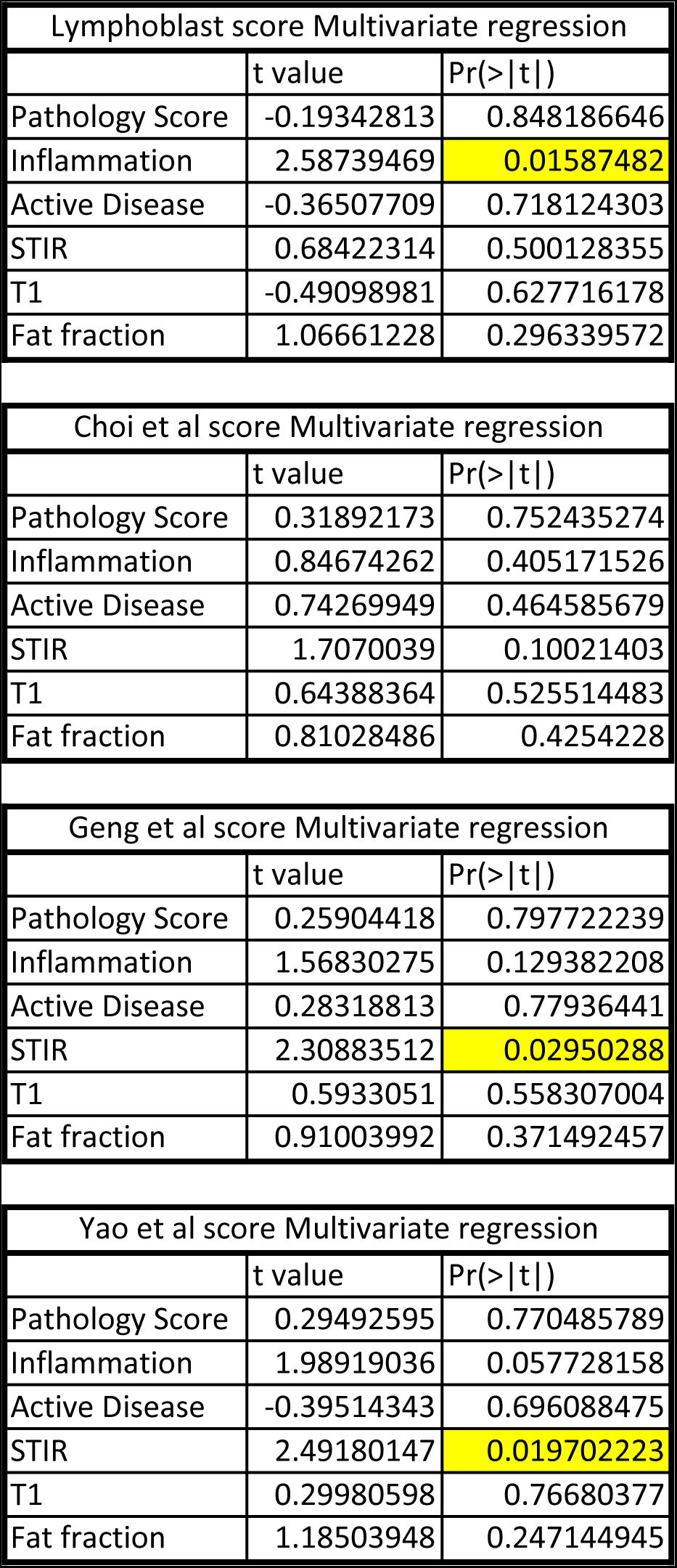
The FSHD Lymphoblast score correlates specifically with histological inflammation in FSHD patient muscle biopsies, while the later DUX4 target gene expression scores correlate with STIR positivity on MRI. Tables summarise multivariate regression analyses on the dataset described by Wang et al., 2019^26^ determining the independent association of histological (Pathology score, Inflammation, Active Disease) and MRI based (STIR, T1, fat fraction) assessments of FSHD disease activity with the FSHD Lymphoblast score and each of the 3 DUX4 target gene signatures in turn. Multivariate regression *t*-values and associated *p*-values are provided for each of the FSHD disease activity variables association with each score separately, *p*-values attaining significance at the 5% level are highlighted in yellow. We see that the FSHD Lymphoblast score is only independently associated with histological inflammation, while the Choi et al., 2016^38^ early DUX4 target signature is not independently associated with any measure of disease activity, and the two late DUX4 target gene signatures (Yao et al., 2014^21^ and Geng et al., 2012^20^) both associate with the level of STIR positivity.

## Discussion

Here we have demonstrated that FSHD patient-derived LCLs express DUX4, as well as early and late DUX4 target genes, more robustly than FSHD patient-derived myoblasts and in vitro differentiated myotubes. We further demonstrate that a set of genes up-regulated in FSHD LCLs are elevated on FSHD muscle biopsies, where they correlate with the level of DUX4 target gene activation, but critically are unaltered on isolated FSHD myoblasts or myotubes. Finally, we show that this FSHD Lymphoblast score specifically associates with the level of microscopic histological inflammation in FSHD muscle biopsies. Taken together, our results indicate that immune cell infiltrates contribute to DUX4 target gene expression in FSHD muscle biopsies, and so it is not solely driven by muscle cells.

FSHD is an enigmatic pathology, characterised by considerable heterogeneity and complex molecular pathophysiology. Despite this, consensus is beginning to emerge on certain aspects^2^ particularly the causal role of DUX4 in driving FSHD: a theory underpinned by the genetic basis of the condition^13–15^. However, understanding how DUX4 causes pathology has proven difficult. FSHD presents as a myopathy, hence studies into DUX4 in FSHD have typically focused on muscle cells^19–22,38^. DUX4 is very difficult to detect in FSHD muscle tissue though, generally requiring techniques such as nested RT-qPCR, with immunolabelling detecting DUX4 in as few as 1/1000 proliferating FSHD myoblasts^22,44^. Indeed, by RNA-seq we were unable to detect a single DUX4 transcript in 18 FSHD myoblast samples.

While investigation of FSHD muscle cells is important, muscle is not a homogenous tissue, and given the genetic basis of FSHD, it is likely that non-muscle cell types also express DUX4. The pathological muscle damage observed in FSHD may not solely be driven by deficits in muscle, but also by aberrant inflammation and vascularisation. Indeed, FSHD muscle biopsies are characterised by lymphocytic infiltrates, particularly of endomysial and perivascular CD8^+^ T lymphocytes^27^, while capillary density is significantly lower^47^. Here we show that FSHD patient derived lymphocyte cell lines, in contrast to muscle cell lines, express DUX4 and DUX4 target genes in a robust manner. Furthermore, genes up-regulated in FSHD LCLs are also up-regulated in FSHD muscle biopsies, where they associate strongly with histological assessment of inflammatory infiltrates. Importantly, these LCL genes are unaltered in isolated FSHD myoblasts and myotubes. Lastly, we show that our FSHD Lymphoblast score correlates with both early and late DUX4 target gene expression in FSHD muscle biopsies. These results indicate that FSHD lymphocytes express DUX4 and DUX4 targets at constitutive, high levels and that the characteristic lymphocytic infiltration in FSHD muscle biopsies likely contributes to DUX4 target gene expression in such samples.

A muscle cell contribution to DUX4 in FSHD muscle biopsies is also likely, but via dynamic, stochastic DUX4 expression rather than the continuous expression seen in lymphocytes^32,40^. Time points for these transient bursts of DUX4 expression could include in satellite cells, when DUX4 expression may interfere with the normal function of the transcription factor PAX7^19,23,24^, and during myogenic differentiation, where DUX4 target gene expression may interfere with normal myogenesis^32,40,48^. Here we provide evidence for transient DUX4 expression at both these time points. We demonstrate that while FSHD myoblasts lack expression of both DUX4 and early DUX4 target genes, they exhibit clear up-regulation of later DUX4 target genes – indicating a historic, transient expression of DUX4. Such expression in satellite cell-derived myoblasts could explain the robust repression of PAX7 target genes seen in FSHD muscle biopsies^23,24^. We further demonstrate that DUX4 is detectible by RNA-seq in 17% of FSHD myotube samples, which also display clear up-regulation of both early and late DUX4 target genes compared to their corresponding myoblast samples. This is in line with a transient up-regulation of DUX4 and its target genes during myogenesis, a finding supported by a burst-like expression pattern of DUX4 we reveal in RNA-seq of single FSHD unfused myocytes. Such dynamic DUX4 up-regulation may contribute to the modest efficacy of DUX4 target gene expression as a biomarker in FSHD muscle biopsies^23^, but this is likely in combination with contributions from DUX4 expressing immune cells.

Our findings have a number of implications. The first relates to DUX4 function and role in pathology. Currently, investigation of DUX4 target genes in FSHD has been performed in myoblast cell lines^18,20,21,38,41^. These target genes lead to pro-apoptotic and anti-myogenic effects^18,38^ but importantly, certain immune system processes have also been found dysregulated by DUX4^20^. In addition, a DUX4-IGH fusion gene is present in a significant proportion of adult B-ALL, where it binds DUX4 response elements, and inhibits lymphocytic differentiation^34,35^. These findings point to a role for DUX4 in modification of immune cell function. Here we show that DUX4 is continuously expressed in FSHD LCLs and that DUX4 target genes derived from myoblast over-expression studies, are robustly expressed in FSHD LCLs. This suggests a ubiquity in DUX4 target gene activation regardless of cell type, indicating that processes altered in myogenic cells in FSHD may similarly be altered in FSHD immune cells. As DUX4 and its target genes likely contribute to FSHD progression and are targets for potential therapeutics^49–51^, this suggests a role for the immune system, as well as muscle cells, in contributing to DUX4-mediated pathology in FSHD.

Histology and MRI have long pointed to a role for inflammation in contributing to FSHD muscle damage^26,27,46^, and here we show that a signature derived from circulating immune cells in FSHD patients, correlates with this inflammation. Importantly, the FSHD Lymphoblast score associated specifically with early, microscopic histological inflammation in FSHD muscle biopsies while the late DUX4 target genes, the subject of recent biomarker investigations^21,26,32^, associated with later macroscopic inflammation, as assessed by STIR positivity on MRI. This suggests that the FSHD Lymphoblast score may be superior to late DUX4 target gene expression biomarkers in detection of early FSHD pathological inflammation, at possibly reversible stage.

Although anti-inflammatory agents such as corticosteroids have been used in clinical trials for FSHD without obvious benefit, the premise was that the inflammation was in response to muscle pathology and effects on long term disease progression were not assessed^52^. Moreover, expression of DUX4 and its target genes in lymphocytes may alter their function to make them directly pathogenic. As such, if lymphocytes are a primary driver of FSHD, rather than a secondary response, they may require more bespoke therapeutic interventions.

Given that the FSHD Lymphoblast score was derived from circulating blood borne lymphocytes, this raises the possibility of blood cell-based biomarkers for FSHD, to measure severity and progression. This minimally invasive method means that samples can be regularly obtained, removing the need for repeated, often painful, muscle biopsies for disease progression in routine clinical follow up and in FSHD clinical trials. Though blood-based biomarkers for FSHD have been investigated previously, these have been based on panels of serological tests, rather than assessment of blood borne cell transcriptomes and thus cannot be expected to link to the FSHD causal DUX4 gene and its targets^53,54^. Importantly, our blood cell-derived biomarker demonstrated discriminatory power and correlation with disease activity when assessed on FSHD muscle biopsy transcriptomic samples. This suggests that gene expression changes in patient lymphocytes may reflect changes in the pathologically damaged muscle. Such a finding requires confirmation with paired blood samples and muscle biopsies, beyond the scope of this proof-of-principle study. Moreover, many putative FSHD therapeutics in development target DUX4 and target gene expression^49,51,55,56^. We have shown that DUX4 and its target genes are robustly expressed in FSHD blood cells in a manner correlated with pathological muscle changes, thus anti-DUX4 therapy may be easily monitored for dose-titration by blood cell based DUX4 expression assessment.

To summarise, we demonstrate that FSHD blood borne lymphocytes continuously express DUX4 and its target genes, in contrast to a burst like pattern seen in muscle cells. FSHD lymphocyte gene expression also contributes to DUX4 target gene expression in FSHD muscle biopsies, and associates with early stage microscopic histological inflammation. We propose that FSHD immune cells may contribute to DUX4-mediated pathology in FSHD, and that blood cell based transcriptomic biomarkers may prove useful minimally-invasive new tools for the assessment of FSHD disease activity, progression and therapeutic monitoring.

## Materials and Methods

### Cell Culture of FSHD LCLs and primary myoblasts

Lymphoblastoid FSHD cell lines GSM16283, GSM16414, GSM16278 and respective control lines GSM16281, GSM16320, GSM16412 were cultured in suspension in RPMI-1640 medium, supplemented with L-glutamine, sodium bicarbonate (Sigma), 10% FBS (Sigma) and gentamycin (Gibco). Cell pellets were collected from three independent flasks for each cell line.

Cell pellets corresponding to FSHD primary myoblast cell lines FSHD3 (FSHD1, 7RU, female), FSHD6 (FSHD1, 8RU, female) and FSHD9 (FSHD1, 7RU, male) alongside age and gender matched controls, in proliferation and after 3 days of differentiation into multinucleated myotubes, in singlet, were kind gifts from Dr Dalila Laoudj-Chenivesse.

### RNA-sequencing of FSHD LCLs and primary myoblasts

RNA was isolated using miRNeasy kit (Qiagen) including a DNAse digestion step. Prior to sequencing RNA was analysed by LabChip Bioanalyzer, Qubit fluoremetric quantification and Nanodrop quantification concentration and stability. RNA-seq libraries were prepared using the sureselect stranded RNAseq protocol (Illumina), which allows polyA selection but was modified to work with ribodepletion (Agilent). Libraries were sequenced on an Illumina HiSeq2500.

Raw reads were trimmed using trim-galore, utilising cutadapt14 (v0.4.0) to remove the Illumina Sequencing Adapter (AGATCGGAAGAGC) at the 3’ end. Additionally, 12 bases were also trimmed from the 5’ end, in both myoblast and LCL samples and 5 bases from the 3’ end in the LCL samples, since they showed a biased distribution. Reads were mapped to the human transcriptome using the human genome sequence GRCh38 and v82 gene annotations downloaded from Ensembl. Mapping was performed using tophat 15 (v2.1.0) and bowtie 16 (v1.1.0), enabling the fr-firststrand option of tophat to restrict mapping to the sense strand of the transcript. Reads were assigned to genes using the featureCounts program 17 (v1.5.0), counting fragments and ignoring multi-mapping reads, and restricted to the sense strand. The resulting matrix of read counts was analysed using R.

Data describing the myoblast and LCLs were processed in separate batches and therefore analysed as separate datasets. Both datasets were normalised using the DESeq2 package^57^ in R.

### Public data on FSHD myoblasts, myotubes and muscle biopsies

Data containing myoblast and myotube RNA-seq samples in triplicate from immortalised FSHD myoblast cell lines 54-2, 54-12, 54-A5, 16ABic and 16UBic and matched controls 54-A10, 54-6, 16UBic and 12UBic that we previously described^23,36^ are available from the GEO database, accession numbers: GSE123468 and GSE102812. This data describes 27 (18 FSHD, 12 control) myoblast samples and 27 (18 FSHD, 12 control) myotube samples.

Data containing RNA-seq of 7234 (5133 FSHD, 2101 control) single myocytes was described by van den Heuvel et al., 2019^32^, and normalised read counts were downloaded from GEO database accession GSE122873.

Seven data sets containing transcriptomic assessments of muscle biopsies were analysed, all were downloaded as normalised data sets from the GEO database. Rahimov et al., 2012^42^, GSE36398, describes 50 muscle biopsies assessed by microarray. Bakay et al., 2006^45^, GSE3307, describes 30 muscle biopsies assessed by microarray. Tasca et al., 2012, GSE26852, describes 15 muscle biopsies assessed by microarray. Osborne et al., 2007, GSE10760, describes 49 muscle biopsies assessed by microarray. Dixit et al., 2007, GSE9397, describes 18 muscle biopsies assessed by microarray. Yao et al., 2014, GSE56787, describes 23 muscle biopsies assessed by RNA-seq (control sample C6 was removed as it was the only non-quadriceps sample). Wang et al., 2019, GSE115650, describes 43 muscle biopsies assessed by RNA-seq. Together these 7 datasets describe 228 muscle biopsies (130 FSHD, 98 control).

All data was log-transformed and quantile normalised within study for computation of the DUX4, FSHD Lymphoblast and PAX7 scores, in line with our previously described methodology^23^.

### DUX4 detection, differential expression analysis and derivation of the FSHD Lymphoblast Score

In all RNA-seq datasets, DUX4 detection was reported as positive if a single read was present in the normalised data set.

Differential expression analysis of the LCL data was performed using the DESeq2 package in R^57^ to identify genes associated with FSHD independently of gender and matched-control pair, feature significance was confirmed via *p*-value histogram.

The top 500 significant genes were considered for further analysis. The FSHD LCLs were found to express high levels of DUX4 and DUX4 target genes, and DUX4 is a transcriptional activator with repressed genes adding no power in previous FSHD biomarkers^23^. We thus considered the mean expression of the 237/500 genes that were upregulated in FSHD LCLs in a given sample, as a potential FSHD biomarker, referred to as the FSHD Lymphoblast score.

### Statistics

Biomarker computation and evaluation: Computation of the three DUX4 expression biomarkers and PAX7 target gene repression biomarker were as previously described^23,24^. Briefly, each DUX4 target gene expression score is computed for each sample as the mean expression of the genes found to be up-regulated by the studies of Yao et al., 2014^21^ (114 genes), Geng et al., 2012^20^ (165 genes) and Choi et al., 2016^38^ (212 genes). The PAX7 target gene repression score for each sample was computed as the *t*-score from a test comparing the up-regulated (311 genes) to down-regulated (290 genes) PAX7 target genes within each sample. We have published a software for the computation of each of these scores from suitably normalised dataset^24^. The FSHD Lymphoblast score was computed in each sample as the mean expression of the 237 genes found up-regulated in FSHD LCLs.

For myoblast, myotube and LCL samples, the 3 DUX4 scores and the FSHD Lymphoblast score were evaluated and *z*-normalised within matched control pairs. Score differences between FSHD and controls samples were then evaluated within each cell type via a Wilcoxon *U*-test. ROC curve analysis and AUC computation was performed using the pROC package in R^58^.

For FSHD muscle biopsy samples the 3 DUX4 scores, the FSHD Lymphoblast score and the PAX7 score were computed for each sample and *z*-normalised within each of the 7 studies. Score differences between FSHD and control samples were evaluated within each study via Wilcoxon *U*-test. In the case of the FSHD Lymphoblast score, meta-analysis across the 7 independent studies was performed using a random effects model, and overall significance assessed via Fisher’s combined test. ROC curve analysis, AUC computation and DeLong’s test were performed using all *z*-normalised scores for all studies combined, via the pROC package in R^58^.

Correlation analyses: Pearson correlations between the 3 DUX4 scores and DUX4 expression were computed using the base package in R separately across LCL, myoblast, myotube and single cell myocyte samples following *z*-normalisation within control matched pairs. Pearson correlations between the 3 DUX4 scores and the FSHD Lymphoblast score were computed using the base package in R, following *z*-normalisation within each of the 7 studies considered.

In the case of the muscle biopsy dataset described by Wang et al., 2019^26^, a multivariate regression model was built for the FSHD Lymphoblast score and each of the 3 DUX4 scores to assess independent associations with the 3 histopathological and 3 MRI based measures of disease activity paired with the RNA-seq samples.

### Study approval

Lymphocyte lines used were described in Jones et al., 2017^33^, where ethical permission is detailed. Primary FSHD and control myoblasts were described in Barro et al. 2010^16^ where ethical permission is detailed.

## Author contributions

CSRB: designed research study, conducted experiments, acquired data, analyzed data, and wrote the manuscript.

MP: conducted experiments, acquired data.

PSZ: designed research study and wrote the manuscript.

## Acknowledgements

FSH Society (FSHS-82016-03 to C.R.S.B. and P.S.Z.); Foulkes Foundation Fellowship (to C.R.S.B.); M.P. was funded by an FSH Society fellowship (FSHS-82017-05). The Zammit laboratory is supported in this project by the Medical Research Council (MR/P023215/1); FSH Society Shack Family and Friends researchgrant (FSHS-82013-06) and Association Française contre les Myopathies (AFM 17865).

## Supplemental Information

**Figure S1:**
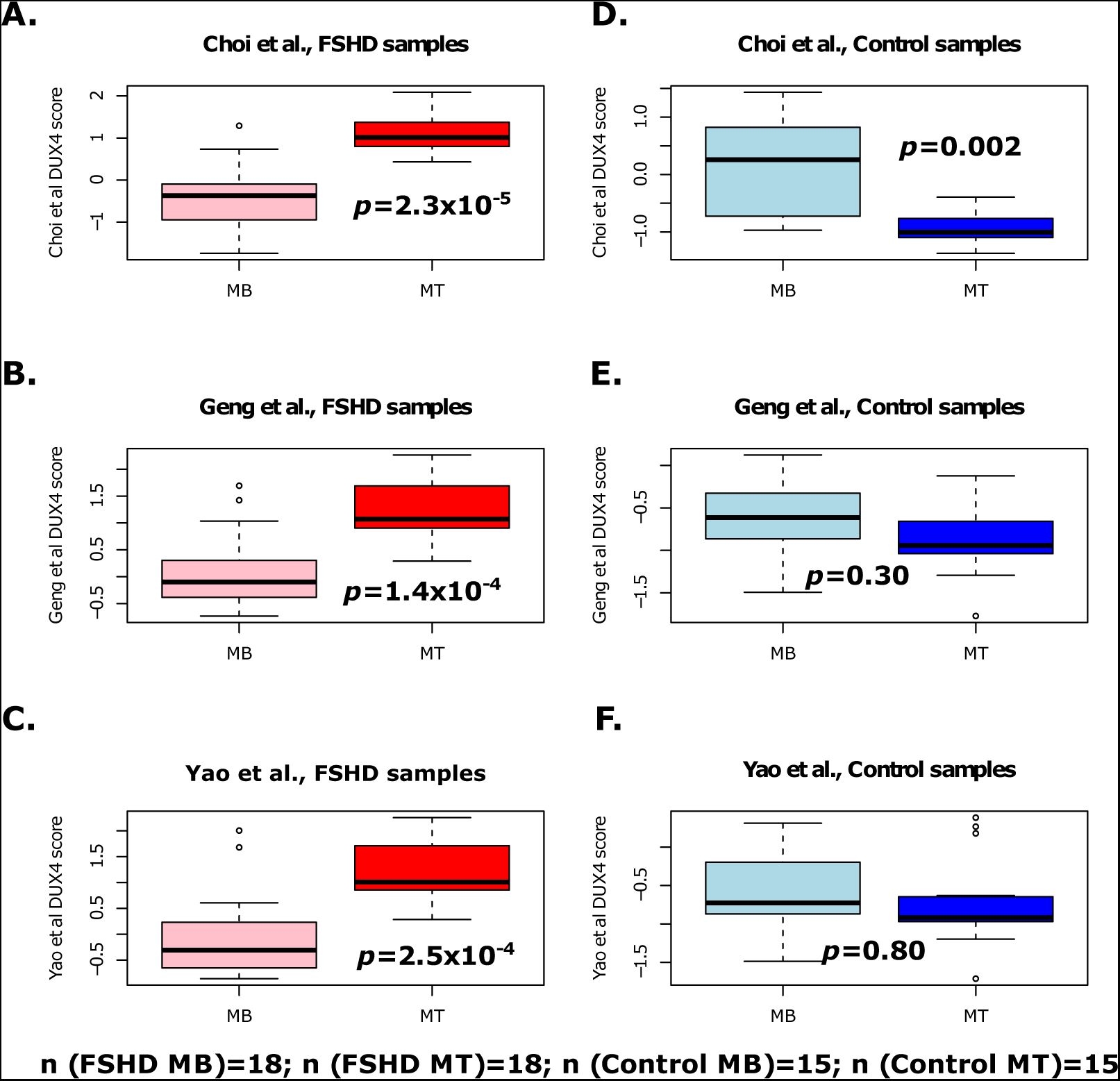
FSHD myotubes show upregulation of both early and late DUX4 target genes compared to matched myoblasts. Boxplots demonstrate the Choi et al., 2016^38^ early (8 hour) DUX4 target gene signature, the Yao et al., 2014^21^ (24-48 hour) and Geng et al., 2012^20^ later (24 hour) DUX4 target gene signature (*z-*normalised within FSHD patient matched control groups) in myoblasts and myotubes from FSHD samples (A-C) and control samples (D-F). The box represents the interquartile range (IQR), with the median indicated by a line. Whiskers denote min (1.5*IQR, max (observed value)). ‘‘o’’ represents data points greater than 1.5 IQR from the median. Wilcoxon *U*-test *p*-values comparing myoblast to myotube samples are presented on each plot. In FSHD samples, the 3 DUX4 target gene signatures are elevated on myotubes compared to matched myoblast samples. In control samples, the Choi et al., 2016^38^ early DUX4 target gene signature is suppressed in myotubes compared to myoblasts, while the two later DUX4 target gene signatures are unaltered.

**Figure S2:**
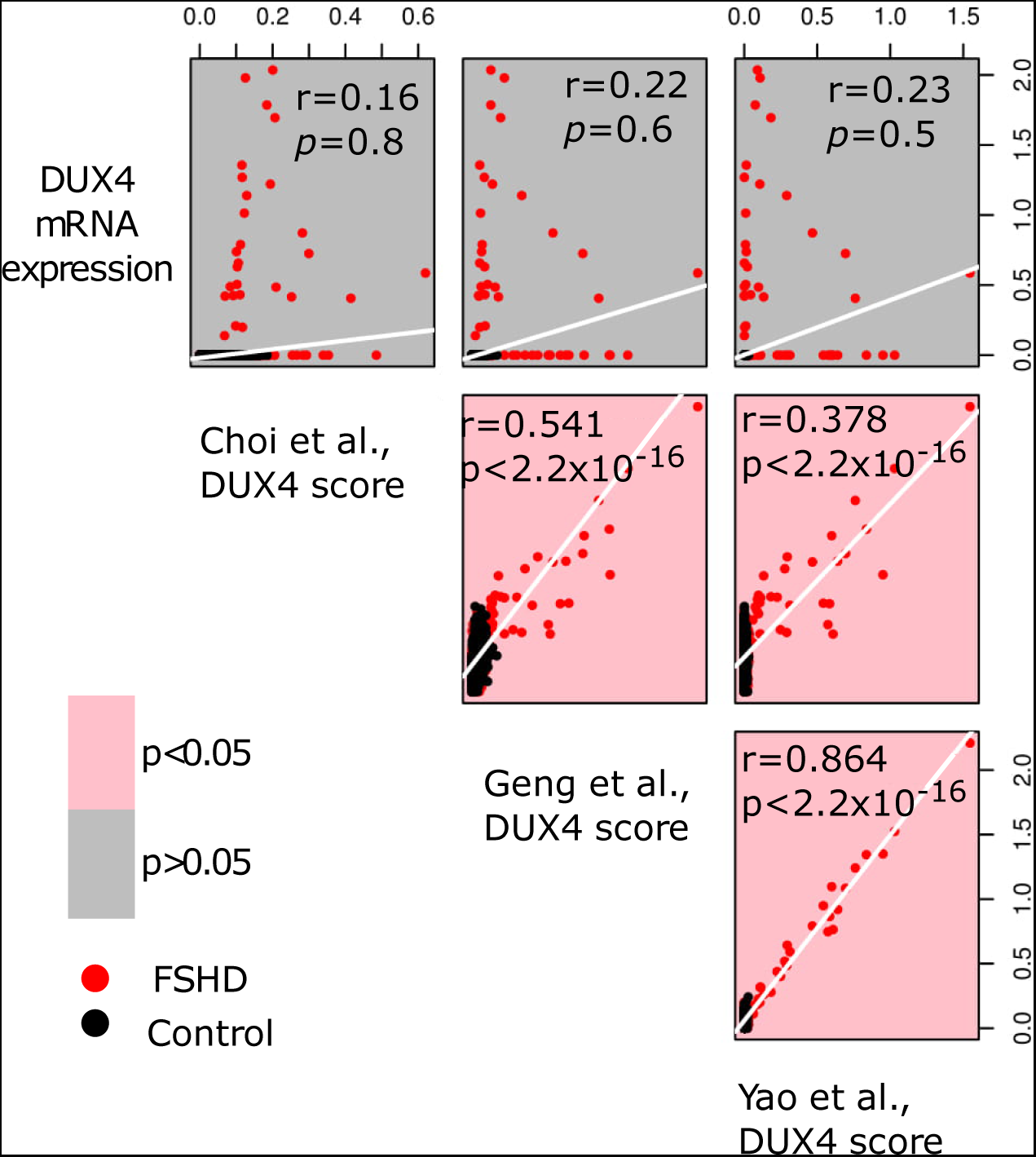
DUX4 displays a burst like expression pattern in single cell RNA-seq of FSHD patient myocytes. Scatter plots display DUX4 expression, the Choi et al., 2016^38^ early (8 hour) DUX4 target gene signature, and the Yao et al., 2014^21^ later (24-48 hour) and Geng et al., 2012^20^ later (24 hour) DUX4 target gene signatures, plotted against one another across the 7234 single myocyte samples from 2 FSHD1, 2 FSHD2 and 2 control patients in the dataset described by van den Heuvel et al., 2019^32^. Pearson’s *r* and associated *p*-value is provided for each pairwise comparison, in the case of DUX4 mRNA, correlations are only made over the 27 DUX4 expressing cells. Red points correspond to FSHD samples and black points correspond to controls. Plots denoting correlations reaching significance are pink, whilst those not attaining significance are grey. DUX4 mRNA expression follows a burst like expression pattern, attaining high values whilst DUX4 target gene expression is low and dropping as the target gene expression rises

**Figure S3:**
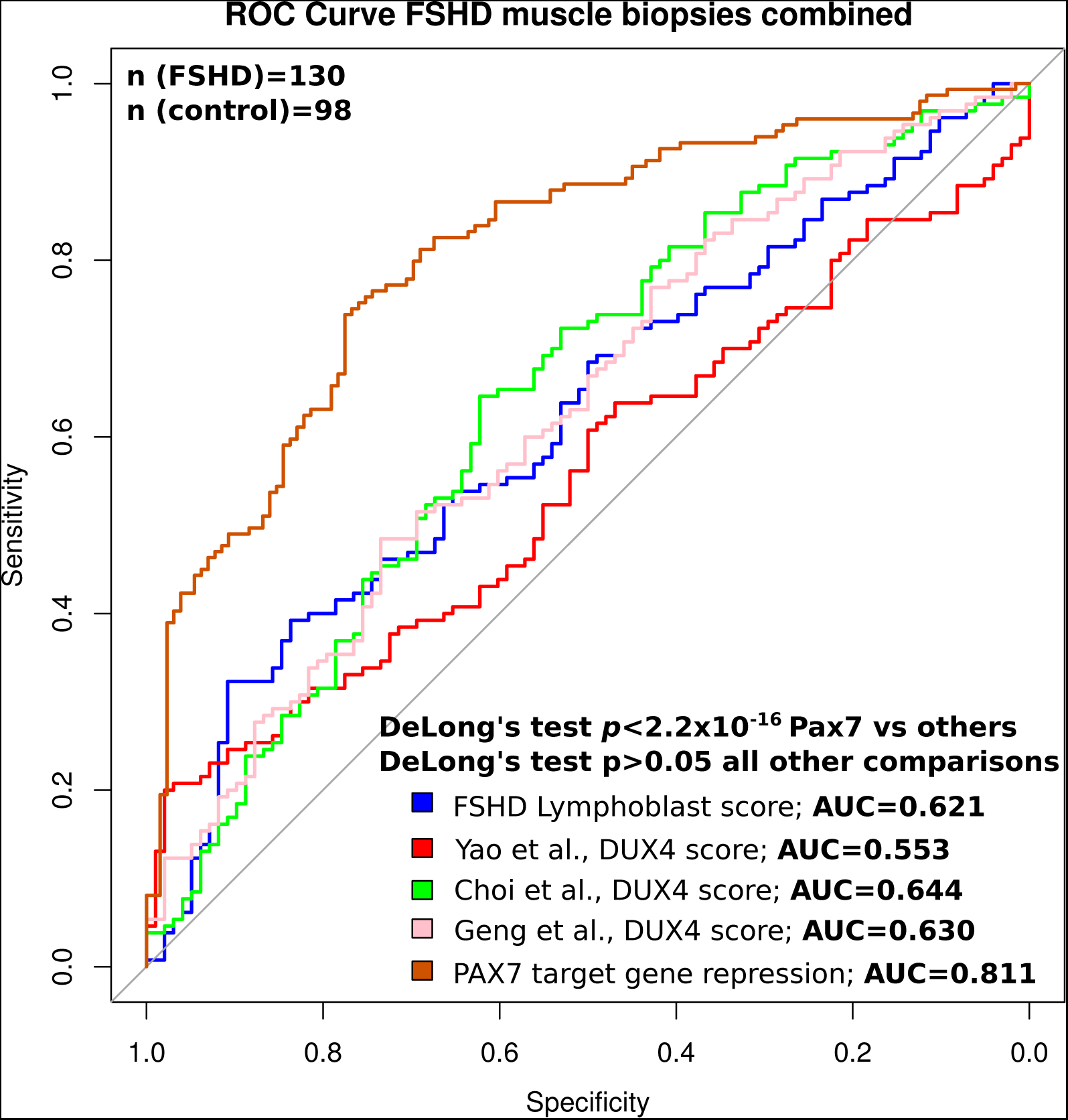
The FSHD Lymphoblast score represents a moderately powered biomarker for FSHD muscle biopsies, equivalent to DUX4 target gene expression but inferior to PAX7 target gene repression. A ROC curve displays the discriminatory capacity of the FSHD Lymphoblast score (blue), the Choi et al., 2016^38^ early (8 hour) DUX4 target gene signature (green), the Yao et al., 2014^21^ later (24-48 hour) DUX4 target gene signature (red), the Geng et al., 2012^20^ later (24 hour) DUX4 target gene signature (pink) and the PAX7 target gene repression score (brown) on all muscle biopsy datasets combined. Each score was computed on each muscle biopsy sample and *z*-normalised within each of the 7 independent studies before being pooled for ROC curve analysis. The AUC for each score is displayed alongside DeLong’s test *p*-value comparing the discriminatory power of the biomarkers. FSHD Lymphoblast score displays a discriminatory power similar to DUX4 target gene signatures, but inferior to PAX7 target gene repression.

**Table S1: FSHD LCL Score Genes**

## References

1. Deenen, J. C. W. et al. Population-based incidence and prevalence of facioscapulohumeral dystrophy. Neurology 83, 1056–1059 (2014).

2. Tawil, R., van der Maarel, S. M. & Tapscott, S. J. Facioscapulohumeral dystrophy: the path to consensus on pathophysiology. Skelet. Muscle 4, 12 (2014).

3. Orrell, R. W. Facioscapulohumeral dystrophy and scapuloperoneal syndromes. in Handbook of clinical neurology 101, 167–180 (2011).

4. Sacconi, S., Salviati, L. & Desnuelle, C. Facioscapulohumeral muscular dystrophy. Biochim. Biophys. Acta 1852, 607–14 (2015).

5. Tawil, R., Storvick, D., Feasby, T. E., Weiffenbach, B. & Griggs, R. C. Extreme variability of expression in monozygotic twins with FSH muscular dystrophy. Neurology 43, 345–348 (1993).

6. Sakellariou, P. et al. Mutation spectrum and phenotypic manifestation in FSHD Greek patients. Neuromuscul. Disord. 22, 339–349 (2012).

7. Nikolic, A. et al. Clinical expression of facioscapulohumeral muscular dystrophy in carriers of 1–3 D4Z4 reduced alleles: experience of the FSHD Italian National Registry. BMJ Open 6, e007798 (2016).

8. Fitzsimons, R. B. Retinal vascular disease and the pathogenesis of facioscapulohumeral muscular dystrophy. A signalling message from Wnt? Neuromuscul. Disord. 21, 263–271 (2011).

9. Longmuir, S. Q. et al. Retinal arterial but not venous tortuosity correlates with facioscapulohumeral muscular dystrophy severity. J. Am. Assoc. Pediatr. Ophthalmol. Strabismus 14, 240–243 (2010).

10. Fitzsimons, R. B., Gurwin, E. B. & Bird, A. C. RETINAL VASCULAR ABNORMALITIES IN FACIOSCAPULOHUMERAL MUSCULAR DYSTROPHY A GENERAL ASSOCIATION WITH GENETIC AND THERAPEUTIC IMPLICATIONS. Brain 110 (1987).

11. Lutz, K. L., Holte, L., Kliethermes, S. A., Stephan, C. & Mathews, K. D. Clinical and genetic features of hearing loss in facioscapulohumeral muscular dystrophy. Neurology 81, 1374–1377 (2013).

12. Trevisan, C. Pietro et al. Facioscapulohumeral muscular dystrophy and occurrence of heart arrhythmia. Eur. Neurol. 56, 1–5 (2006).

13. Lemmers, R. J. L. F. et al. A unifying genetic model for facioscapulohumeral muscular dystrophy. Science 329, 1650–1653 (2010).

14. Lemmers, R. J. L. F. et al. Digenic inheritance of an SMCHD1 mutation and an FSHD-permissive D4Z4 allele causes facioscapulohumeral muscular dystrophy type 2. Nat. Genet. 44, 1370–1374 (2012).

15. van den Boogaard, M. L. et al. Mutations in DNMT3B Modify Epigenetic Repression of the D4Z4 Repeat and the Penetrance of Facioscapulohumeral Dystrophy. Am. J. Hum. Genet. 98, 1020–1029 (2016).

16. Barro, M. et al. Myoblasts from affected and non-affected FSHD muscles exhibit morphological differentiation defects. J. Cell. Mol. Med. 14, 275–289 (2010).

17. Zeng, W. et al. Specific loss of histone H3 lysine 9 trimethylation and HP1gamma/cohesin binding at D4Z4 repeats is associated with facioscapulohumeral dystrophy (FSHD). PLoS Genet. 5, e1000559 (2009).

18. Knopp, P. et al. DUX4 induces a transcriptome more characteristic of a less-differentiated cell state and inhibits myogenesis. J. Cell Sci. 129, 3816–3831 (2016).

19. Bosnakovski, D. et al. An isogenetic myoblast expression screen identifies DUX4-mediated FSHD-associated molecular pathologies. EMBO J. 27, 2766–2779 (2008).

20. Geng, L. N. et al. DUX4 activates germline genes, retroelements, and immune mediators: implications for facioscapulohumeral dystrophy. Dev. Cell 22, 38–51 (2012).

21. Yao, Z. et al. DUX4-induced gene expression is the major molecular signature in FSHD skeletal muscle. Hum. Mol. Genet. 23, 5342–52 (2014).

22. Snider, L. et al. Facioscapulohumeral Dystrophy: Incomplete Suppression of a Retrotransposed Gene. PLoS Genet. 6, e1001181 (2010).

23. Banerji, C. R. S. et al. PAX7 target genes are globally repressed in facioscapulohumeral muscular dystrophy skeletal muscle. Nat. Commun. 8, 2152 (2017).

24. Banerji, C. R. S. & Zammit, P. S. PAX7 target gene repression is a superior FSHD biomarker than DUX4 target gene activation, associating with pathological severity and identifying FSHD at the single-cell level. Hum. Mol. Genet. (2019).

25. Bosnakovski, D. et al. The DUX4 homeodomains mediate inhibition of myogenesis and are functionally exchangeable with the Pax7 homeodomain. J. Cell Sci. 130, 3685–3697 (2017).

26. Wang, L. H. et al. MRI-informed muscle biopsies correlate MRI with pathology and DUX4 target gene expression in FSHD. Hum. Mol. Genet. (2018). doi:10.1093/hmg/ddy364

27. Frisullo, G. et al. CD8+ T Cells in Facioscapulohumeral Muscular Dystrophy Patients with Inflammatory Features at Muscle MRI. J. Clin. Immunol. 31, 155–166 (2011).

28. Munsat, T. L., Piper, D., Cancilla, P. & Mednick, J. Inflammatory myopathy with facioscapulohumeral distribution. Neurology 22, 335–47 (1972).

29. Figarella-Branger, D., Pellissier, J. F., Serratrice, G., Pouget, J. & Bianco, N. [Immunocytochemical study of the inflammatory forms of facioscapulohumeral myopathies and correlation with other types of myositis]. Ann. Pathol. 9, 100–8 (1989).

30. Arahata, K. et al. Inflammatory response in facioscapulohumeral muscular dystrophy (FSHD): immunocytochemical and genetic analyses. Muscle Nerve. Suppl. 2, S56–66 (1995).

31. Turki, A. et al. Functional muscle impairment in facioscapulohumeral muscular dystrophy is correlated with oxidative stress and mitochondrial dysfunction. Free Radic. Biol. Med. 53, 1068–1079 (2012).

32. van den Heuvel, A. et al. Single-cell RNA-sequencing in facioscapulohumeral muscular dystrophy disease etiology and development. Hum. Mol. Genet. (2018). doi:10.1093/hmg/ddy400

33. Jones, T. I., Himeda, C. L., Perez, D. P. & Jones, P. L. Large family cohorts of lymphoblastoid cells provide a new cellular model for investigating facioscapulohumeral muscular dystrophy. Neuromuscul. Disord. 27, 221–238 (2017).

34. Dong, X. et al. Structural basis of DUX4/IGH-driven transactivation. Leukemia 32, 1466–1476 (2018).

35. Yasuda, T. et al. Recurrent DUX4 fusions in B cell acute lymphoblastic leukemia of adolescents and young adults. Nat. Genet. 48, 569–574 (2016).

36. Banerji, C. R. S. et al. Dynamic transcriptomic analysis reveals suppression of PGC1α/ERRα drives perturbed myogenesis in facioscapulohumeral muscular dystrophy. Hum. Mol. Genet. (2018). doi:10.1093/hmg/ddy405

37. Krom, Y. D. et al. Generation of Isogenic D4Z4 Contracted and Noncontracted Immortal Muscle Cell Clones from a Mosaic Patient. Am. J. Pathol. 181, 1387–1401 (2012).

38. Choi, S. H. et al. DUX4 recruits p300/CBP through its C-terminus and induces global H3K27 acetylation changes. Nucleic Acids Res. 44, 5161–5173 (2016).

39. Young, J. M. et al. DUX4 binding to retroelements creates promoters that are active in FSHD muscle and testis. PLoS Genet. 9, e1003947 (2013).

40. Rickard, A. M., Petek, L. M. & Miller, D. G. Endogenous DUX4 expression in FSHD myotubes is sufficient to cause cell death and disrupts RNA splicing and cell migration pathways. Hum. Mol. Genet. 24, 5901–14 (2015).

41. Jagannathan, S. et al. Model systems of DUX4 expression recapitulate the transcriptional profile of FSHD cells. Hum. Mol. Genet. 25, 4419–4431 (2016).

42. Rahimov, F. et al. Transcriptional profiling in facioscapulohumeral muscular dystrophy to identify candidate biomarkers. Proc. Natl. Acad. Sci. U. S. A. 109, 16234–9 (2012).

43. Dixit, M. et al. DUX4, a candidate gene of facioscapulohumeral muscular dystrophy, encodes a transcriptional activator of PITX1. Proc. Natl. Acad. Sci. 104, 18157–18162 (2007).

44. Osborne, R. J., Welle, S., Venance, S. L., Thornton, C. A. & Tawil, R. Expression profile of FSHD supports a link between retinal vasculopathy and muscular dystrophy. Neurology 68, 569–577 (2007).

45. Bakay, M. et al. Nuclear envelope dystrophies show a transcriptional fingerprint suggesting disruption of Rb–MyoD pathways in muscle regeneration. Brain 129, 996–1013 (2006).

46. Tasca, G. et al. Different Molecular Signatures in Magnetic Resonance Imaging-Staged Facioscapulohumeral Muscular Dystrophy Muscles. PLoS One 7, e38779 (2012).

47. Statland, J. M. et al. Immunohistochemical Characterization of Facioscapulohumeral Muscular Dystrophy Muscle Biopsies. J. Neuromuscul. Dis. 2, 291–299 (2015).

48. Balog, J. et al. Increased DUX4 expression during muscle differentiation correlates with decreased SMCHD1 protein levels at D4Z4. Epigenetics 10, 1133–1142 (2015).

49. Wallace, L. M. et al. RNA Interference Inhibits DUX4-induced Muscle Toxicity In Vivo: Implications for a Targeted FSHD Therapy. Mol. Ther. 20, 1417–1423 (2012).

50. Dmitriev, P. et al. DUX4-induced constitutive DNA damage and oxidative stress contribute to aberrant differentiation of myoblasts from FSHD patients. Free Radic. Biol. Med. 99, 244–258 (2016).

51. Cruz, J. M. et al. Protein kinase A activation inhibits DUX4 gene expression in myotubes from patients with facioscapulohumeral muscular dystrophy.; Tel 871–7818 doi:10.1074/jbc.RA118.002633

52. Tawil, R. et al. A pilot trial of prednisone in facioscapulohumeral muscular dystrophy. FSH-DY Group. Neurology 48, 46–9 (1997).

53. Statland, J., Donlin-Smith, C. M., Tapscott, S. J., van der Maarel, S. & Tawil, R. Multiplex Screen of Serum Biomarkers in Facioscapulohumeral Muscular Dystrophy. J. Neuromuscul. Dis. 1, 181–190 (2014).

54. Petek, L. M. et al. A cross sectional study of two independent cohorts identifies serum biomarkers for facioscapulohumeral muscular dystrophy (FSHD). Neuromuscul. Disord. 26, 405–413 (2016).

55. Bosnakovski, D. et al. High-throughput screening identifies inhibitors of DUX4-induced myoblast toxicity. Skelet. Muscle 4, 4 (2014).

56. Campbell, A. E. et al. BET bromodomain inhibitors and agonists of the beta-2 adrenergic receptor identified in screens for compounds that inhibit DUX4 expression in FSHD muscle cells. Skelet. Muscle 7, 16 (2017).

57. Love, M. I., Huber, W. & Anders, S. Moderated estimation of fold change and dispersion for RNA-seq data with DESeq2. Genome Biol. 15, 550 (2014).

58. Robin, X. et al. pROC: an open-source package for R and S+ to analyze and compare ROC curves. BMC Bioinformatics 12, 77 (2011).

